## Supplementary Figures and Tables for "Involvement of an IgE/Mast cell/B cell amplification loop in abdominal aortic aneurysm progression"

### SUPPLEMENTAL MATERIAL

#### Supplemental Tables

##### Table S1: Patients'characteristics.

| | All (n=57) | Histology (n=25) | FC (n=25) | CM (n=37) |
|---|---|---|---|---|
| **Age (yrs)** | 68 +/- 2 | 70 +/- 2 | 73 +/- 2 | 68 +/- 2 |
| **Male** | 86% | 92% | 92% | 84% |
| **Aneurysm location** | | | | |
| Suprarenal | 2.5% | 5% | 0% | 4% |
| Juxtarenal | 2.5% | 5% | 0% | 0% |
| Subrenal | 59% | 58% | 83% | 57% |
| Subrenal+iliac | 36% | 32% | 17% | 39% |
| **Maximum aortic diameter (cm)** | 61 +/- 2 | 61 +/- 3 | 67 +/- 4 | 63 +/- 3 |
| **Clinical Features** | | | | |
| Diabetes | 5% | 5% | 0% | 7% |
| Hypertension | 81% | 89% | 86% | 87% |
| Hyperlipidaemia | 40% | 47% | 58% | 37% |
| Smoking | 83% | 84% | 86% | 80% |
| **Anti-coagulants** | 15% | 17% | 27% | 15% |

Values are mean +/- SEM or %, for all AAA samples, or samples according to processing (some samples were used for several kind of analysis). FC: flow cytometry analysis (MCs and/or B cells); CM: conditioned medium (IgE concentration and/or MC stimulation).

#### Table S2: NAA healthy donors'characteristics.

| | **All** (n=39) | **Histology** (n=7) | **FC** (n=16) | **CM** (n=24) |
|---|---|---|---|---|
| **Age (yrs)** | 55 +/- 3 | 53 +/- 9 | 55 +/- 5 | 56 +/- 4 |
| **Male** | 61% | 80% | 56% | 61% |
| **Atherosclerotic lesion** | | | | |
| None | 28% | 29% | 31% | 25% |
| Fatty streak | 44% | 43% | 31% | 54% |
| Fibrolipidic | 26% | 29% | 31% | 17% |
| Intraplaque hemorrhage | 3% | 0% | 6% | 4% |

Values are mean +/- SEM or %, for all NAA samples, or samples according to processing (some samples were used for several kind of analysis). FC: flow cytometry analysis (MCs and/or B cells); CM: conditioned medium (IgE concentration and/or mast cell stimulation).

#### Table S3: ApoE-RMB survival and pseudoaneurysm occurrence.

| | PBS (n = 17) | DT (n = 18) | p-value |
|---|---|---|---|
| **Pseudoaneurysm** | n = 7 (47%) | n = 8 (44%) | 0.85 |
| **Death*** | n = 0 (0%†) | n = 2 (25%†) | 0.15 |

*, dead mice presented pseudoaneurysms and died 21 days after the beginning of AngII infusion (7 days after DT treatment). †, % of mice with AAA.

### Supplementary Figures

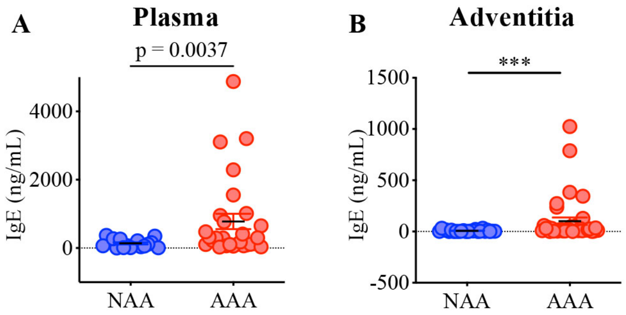


**Fig S1**

#### Fig S1. IgEs are elevated in the plasma and adventitia of AAA patients.

IgEs were titrated in the plasma (A) and conditioned medium from adventitia (B) of NAA organ donors and AAA patients. Mann-Whitney tests were used to compare groups (***: $p < 0.001$).

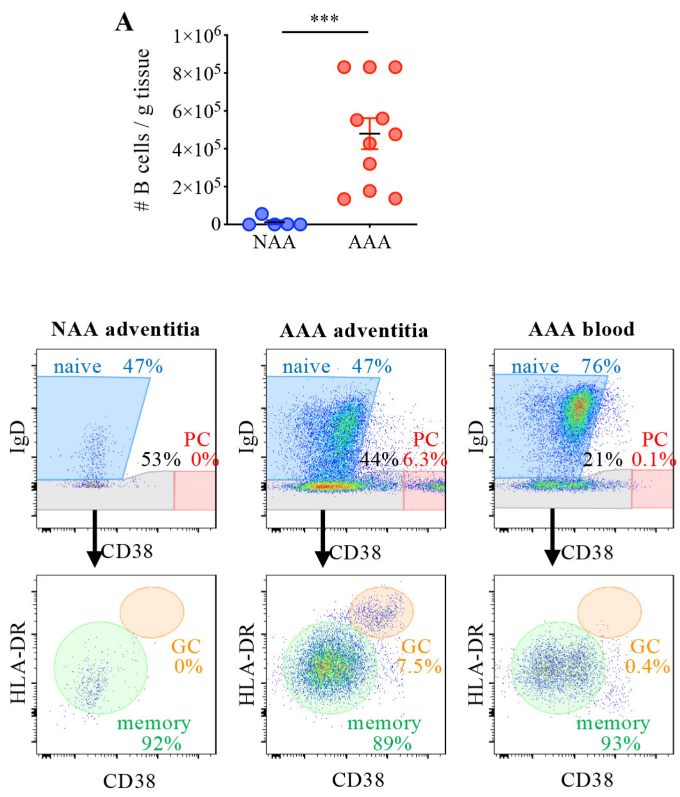


**Fig S2**

#### Fig S2. GC B cells and plasma cells are elevated in AAA adventitia.

Adventitia from NAA organ donors and AAA patients were digested and analysed by flow cytometry after the addition of fluorescent count beads. (A) B cells were identified as singlet, autofluorescent$^-$, live CD45$^+$ CD19$^+$ cells, and their number was calculated in each sample, showing a statistically significant increase of B cells in AAA samples. ***, $p < 0.001$, Mann-Whitney test. (B) B cells were identified as in (A), and subsets were defined as follows: naïve B cells, IgD$^+$ CD38$^-$; plasma cells (PC), IgD$^-$ CD38$^{hi}$; germinal centre B cells (GC), IgD$^-$ CD38$^+$ HLA-DR$^{hi}$; memory B cells, IgD$^-$ CD38$^-$ HLA-DR$^+$. Representative samples show that GC B cells and plasma cells were present in the adventitia of AAA, while they were barely detected in the matched blood of the AAA patient, or in the adventitia of NAA.

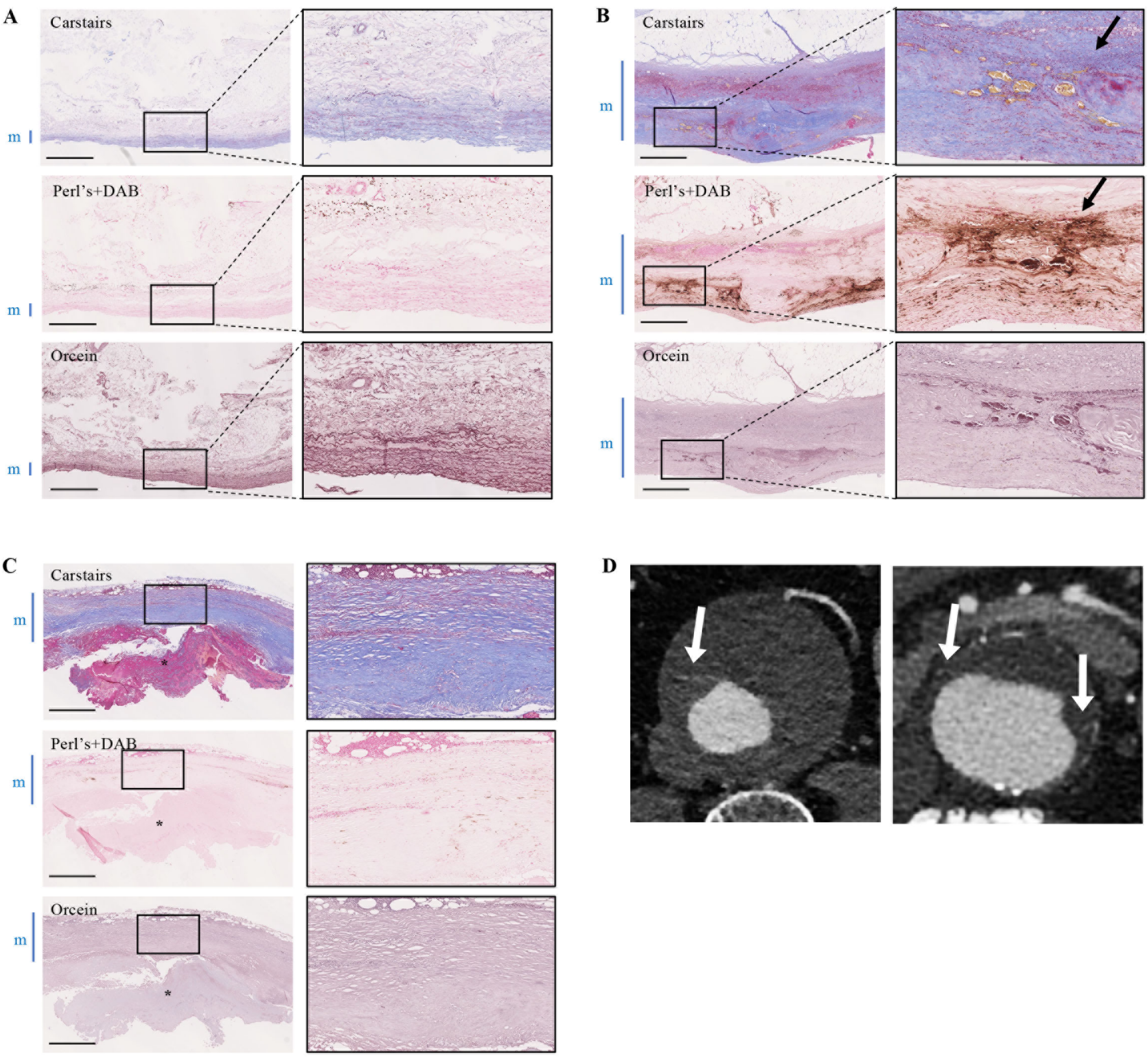


**Fig S3**

#### Fig S3. AAAs with ancient or no intramural haematomas.

Histological staining of NAAs (A) and AAAs (B-C). Serial sections were stained with Carstairs's stain, Perl's+DAB stain and Orcein stain as in Fig 1C. The squares frame the localisation of the details shown in the insets. The lumen is at the bottom of each picture, and the media (m) is indicated by the blue bar on the side of the pictures. In A, intact elastin fibers can be seen in the media on the Orcein stain. In B, black arrows indicate iron deposits in the absence of red blood cells, suggestive of an ancient intramural haematoma. An intraluminal thrombus (*) can be seen in C, where no signs of intramural haematoma were seen. Scale bar: 1 mm. (D) Contrast-enhanced tomography angiograms of AAAs. White arrows: blood disruption from the aortic lumen to the aortic wall through the intraluminal thrombus.

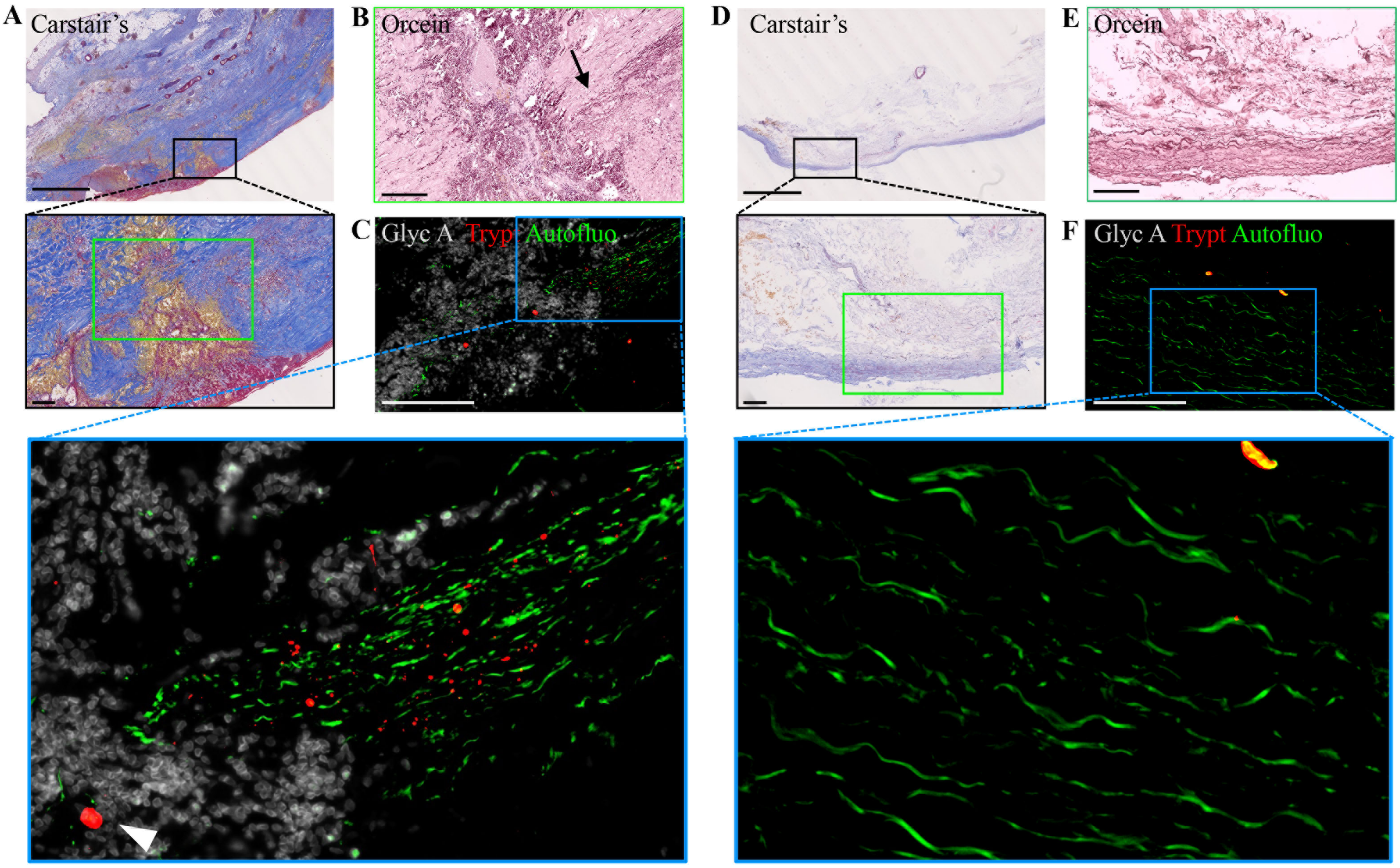


**Fig S4**

#### Fig S4. Proximity of micro-fissures and MC degranulation in human AAAs.

Serial sections of an AAA sample with a micro-fissure (A-C) and from a NAA sample (D-F) were stained with: Carstair's stain (A and D); orcein to reveal elastin fibres (B and E); and tryptase (red) and glycophorin A (gray) to detect MCs/tryptase$^{+}$ granules and red blood cells, respectively (C and F; the elastin fibres were autofluorescent: green). The pictures in B and C, and E and F, correspond to the green insets in A and D, respectively. The entry site of blood into the aortic wall in the AAA sample can be observed on the Carstair's stain (A, red blood cells appear in yellow). The blood entry site visible on the Carstair's stain (A) and with the glycophorin A stain (C) was associated with degraded elastin fibres (black arrow on orcein stain in B) and the presence of MCs (white arrowhead) and tryptase$^{+}$ granules (C). Note the normal aspect of the elastin fibers (E) as well as the absence of tryptase and red blood cells (F) in the control. Scale bars: A and D 2.5 mm; others: 200 µm.

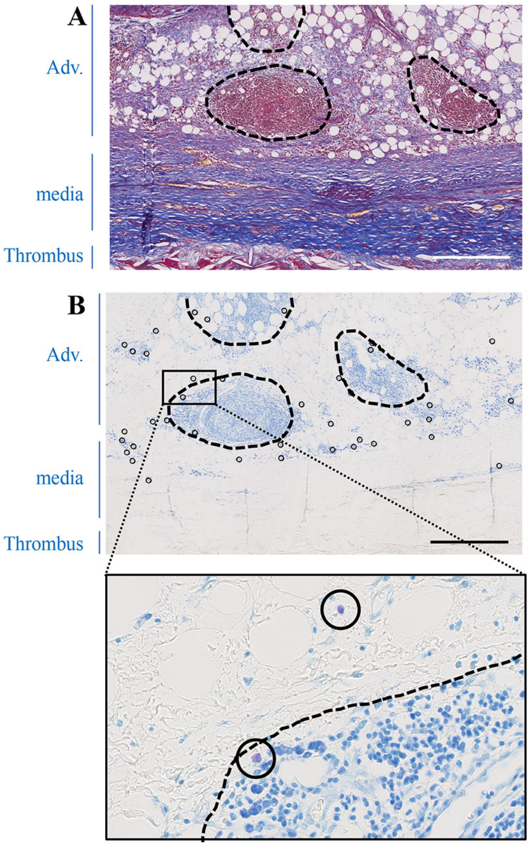


Fig S5

#### Fig S5. MCs accumulate in the adventitia of human AAAs.

Carstairs' stain (A) and toluidine blue stain (B-C) on serial sections of a representative micro-fissured AAA sample. MCs appear purple on toluidine blue stain (indicated with black circles). TLOs are circled with a dotted line. Scale bar: 500 µm.

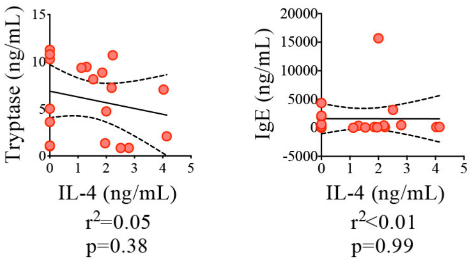


**Fig S6**

#### Fig S6. No correlation between tryptase, IgE and IL-4 in the plasma of AAA patients.

IL-4, tryptase, and IgE concentrations were measured in the serum of AAA patients (same patients as in Fig 3E-F). $r^2$ and p-values from Pearson correlation analysis are indicated.

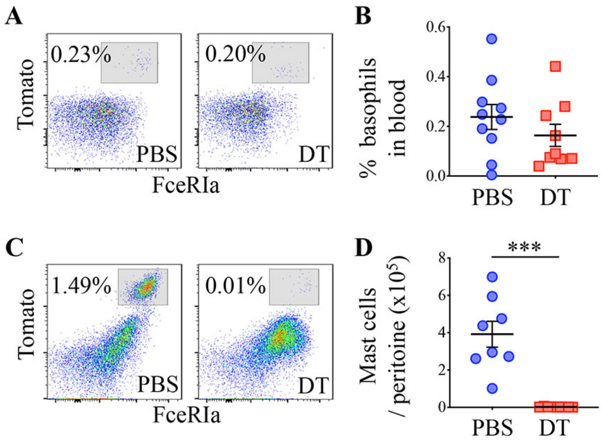


**Fig S7**

#### Fig S7. Repopulation of basophils in the blood and MCs in the peritoneum after DT depletion in ApoE RMB mice.

ApoE-RMB mice were treated as in Fig 4A. At day 14 after DT (n=9) or PBS (n=10) injection (day 28 of Ang II infusion), mice were sacrificed. (A) Blood basophils were identified among singlet cells, as $CD45^+$ Live/$Dead^-$ $CD3^-$ $CD19^-$ $Fc\varepsilon RIa^+$ $Tomato^+$ cells (21). (B) Percentage of basophils within singlet cells, $CD45^+$ Live/$Dead^-$ cells. (C) Peritoneal MCs were identified among singlet cells as $CD45^+$ Live/$Dead^-$ $CD3^-$ $CD19^-$ $Fc\varepsilon RIa^+$ $Tomato^+$ cells $CD117^+$ cells (21). (D) Number of MCs per peritoneal lavage. ***: $p<0.0001$, Mann-Whitney test. Data are representative of two experiments.

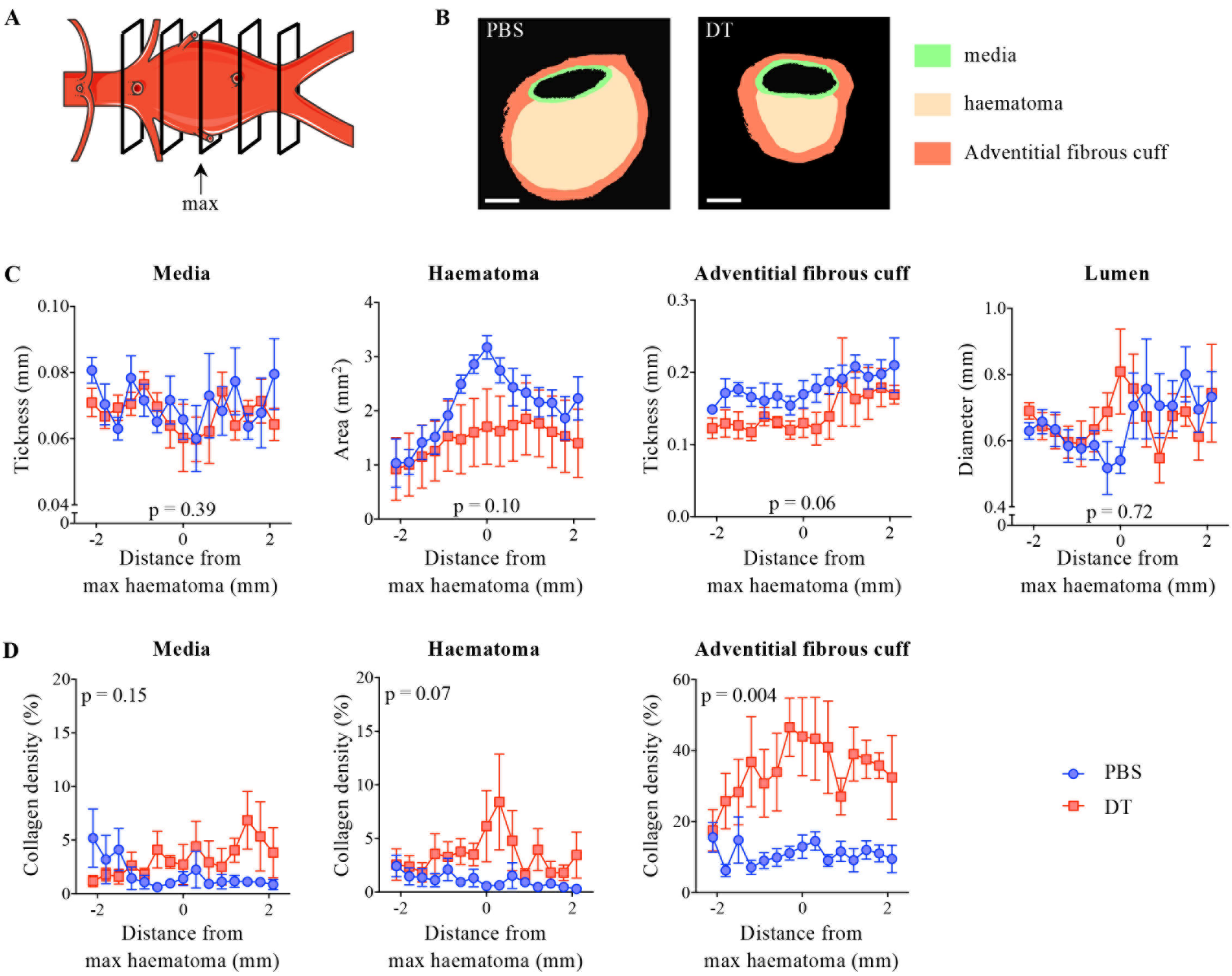


**Fig S8**

#### Fig S8. MCs promote pseudoaneurysm expansion after dissection in ApoE-RMB mice.

(A) Sections taken at different levels (every 300 µm) from each pseudoaneurysm were used for analysis. (B) Vascular wall layers were defined by computer-assisted morphometry on Sirius red-stained cross-sections (Fig 4E) of pseudoaneurysm (scale bar: 500 µm). Size (C) and collagen density (D) of the different layers were calculated by computer-assisted morphometry, and plotted relatively to the distance from the layer with the largest haematoma. Mean +/- standard error; p-values for temperature effect in mixed-model (REML) analysis.
